# Linking the development and functioning of a carnivorous pitcher plant’s microbial digestive community

**DOI:** 10.1101/090225

**Authors:** David W. Armitage

**Affiliations:** Department of Integrative Biology, University of California Berkeley, 3040 Valley Life Sciences Building, Berkeley, CA, USA 94720-3140; Department of Biological Sciences, University of Notre Dame, 100 Galvin Life Science Center, Notre Dame, IN, USA 46556

## Abstract

Ecosystem development theory predicts that successional turnover in community composition can influence ecosystem functioning. However, tests of this theory in natural systems are made difficult by a lack of replicable and tractable model systems. Using the microbial digestive associates of a carnivorous pitcher plant, I tested hypotheses linking host age-driven microbial community development to host functioning. Monitoring the yearlong development of independent microbial digestive communities in two pitcher plant populations revealed a number of trends in community succession matching theoretical predictions. These included mid-successional peaks in bacterial diversity and metabolic substrate use, predictable and parallel successional trajectories among microbial communities, and convergence giving way to divergence in community composition and carbon substrate use. Bacterial composition, biomass, and diversity positively influenced the rate of prey decomposition, which was in turn positively associated with a host leaf’s nitrogen uptake efficiency. Overall digestive performance was greatest during late summer. These results highlight links between community succession and ecosystem functioning and extend succession theory to host-associated microbial communities.

**Statement of authorship:** DWA conceived this work, performed data collection and analysis, and wrote the manuscript

## INTRODUCTION

Although the capacity for community composition to mediate ecosystem processes is widely recognized (Hooper *et al.*, 2005), few theoretical (Finn, 1982; DeAngelis, 1992; Loreau, 1998) and empirical studies (Fisher *et al.*, 1982; Schmidt *et al.*, 2007) have investigated community-ecosystem linkages along natural successional gradients. Ecosystem development theory (Odum, 1969) seeks to explain temporal variation in ecosystem properties in terms of community successional turnover. Central to this theory is the prediction that successional turnover can influence elemental cycling rates leading to a coupling of community composition and ecosystem processes through time (Odum, 1969; Huston and Smith, 1987; DeAngelis, 1992; Loreau, 1998). It is worth noting that modern succession theory does not assume directionality toward a stable equilibrium (or ‘climax’), but instead recognizes that the temporal trajectories of ecosystems can vary due to the relative influences of general ecological processes (Meiners *et al.*, 2015). Although these predictions have not been immune to critique on both theoretical and empirical grounds, adequately replicated tests in natural communities remain scarce.

The natural microcosms of host-associated microbial communities offer a number of unique advantages for testing ecosystem development hypotheses. First, microbiota can enable identifiable and measureable functions for their hosts (Bäckhed *et al.*, 2005; Lugtenberg and Kamilova, 2009). Next, the habitats being colonized are often nearly identical among closely-related individuals, permitting repeated, independent observations of ecosystem development. Finally, the successional dynamics of host-associated microbiota frequently operate over time scales proportional to the host’s lifespan, which can manifest as large shifts in community composition and function over relatively short time periods.

This study uses the microbial digestive communities in developing leaves of the pitcher plant *Darlingtonia californica* (Sarraceniaceae) (Figure 1a) to test the following hypotheses linking community succession to ecosystem function (Figures 1b-e): First, alpha diversity will either asymptotically increase or be unimodal over the host leaf’s lifespan as taxa are recruited from the regional pool and subsequently persist or are excluded by superior competitors (Odum, 1969; Loucks, 1970; Auclair and Goff, 1971; Connell and Slatyer, 1977; Fierer *et al.*, 2010). Consequently, trait diversity (e.g., biochemical pathways, C-substrate use) is also expected to increase as succession proceeds (Odum, 1969). Second, rates of biomass production should decrease over time, as growth-limiting nutrients are lost from the system and/or stored in living biomass — this should manifest as a logistic-like biomass-curve (Odum, 1969; Vitousek and Reiners, 1975; Fierer *et al.*, 2010). Third, beta diversity will increase over time if environmental differences among pitchers cause spatially-variable selection or drift, or decrease over time if different leaves constitute similar selective environments (Christensen and Peet, 1984; Dini-Andreote *et al.*, 2015). Fourth, host ecosystem properties (e.g., nutrient cycling, decomposition) should increase monotonically or be unimodal, concomitant with changes in alpha diversity and biomass, as the accumulation of individuals of different species accelerates the degradation of organic material (Cardinale *et al.*, 2007; Weis *et al.*, 2007; Armitage, 2016). This leads to the prediction that biodiversity and biomass dynamics will set ecosystem processes rates (e.g., decomposition), which, in turn, will set rates on host functioning (e.g., nutrient uptake rates) (Hooper *et al.*, 2005).

**Figure 1.**
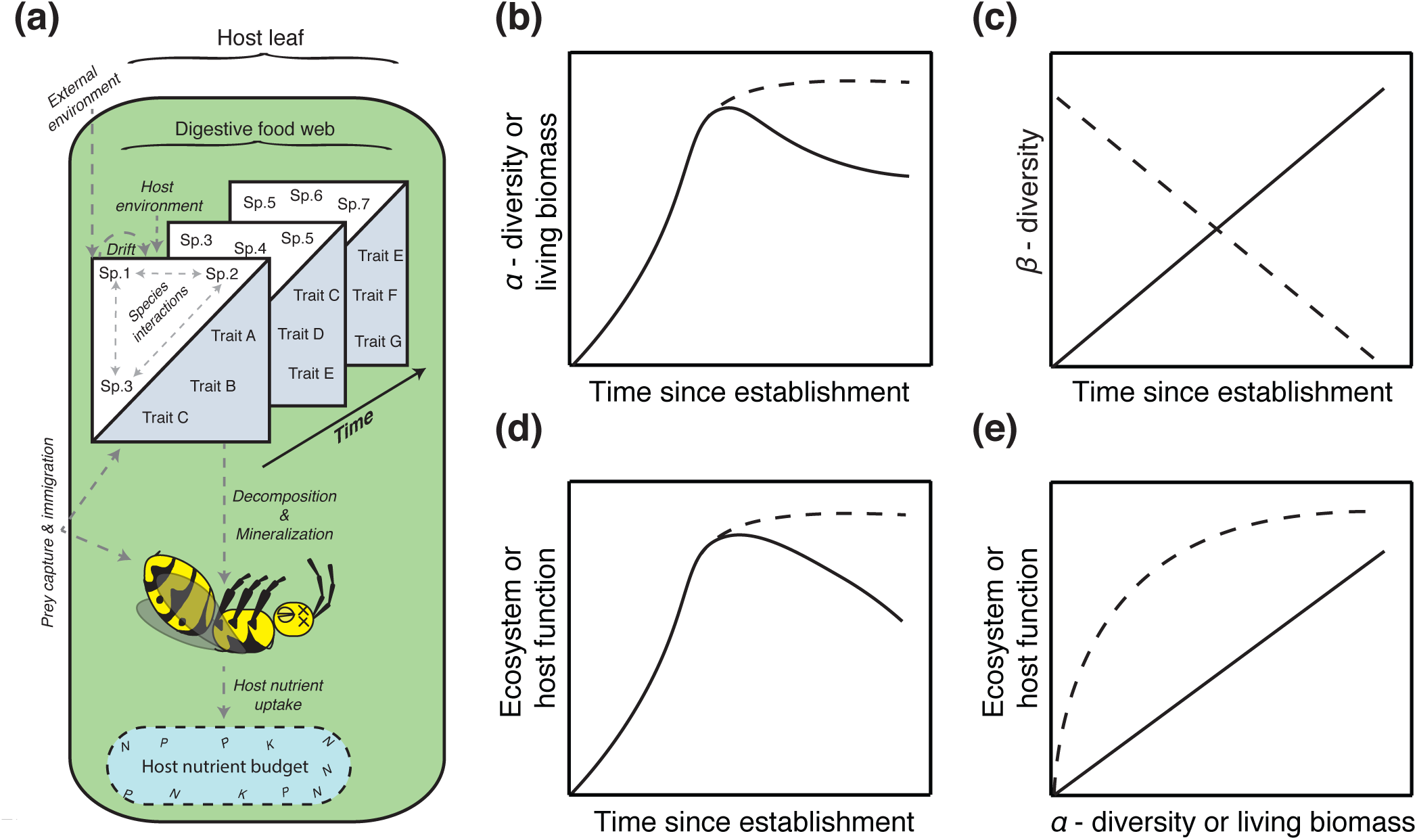
Predictions for successional patterns in *Darlingtonia* leaves. **(a)** Conceptual model for the interactions between host leaf (shaded oval) and its digestive food web (boxes). Dashed grey arrows denote ecological processes hypothesized to influence food web dynamics and host functioning. **(b)** *α*-diversity and living biomass are predicted to increase in pitcher leaves after opening, and eventually either saturate or decrease, consistent with observations made across a variety of ecosystems (Odum, 1969; Loucks, 1970; Auclair and Goff, 1971; Vitousek and Reiners, 1975; Connell and Slatyer, 1977; Peet and Christensen, 1988; Alday *et al.*, 2011). **(c)** Compositional differences among leaf communities (*β*-diversity) may either decrease or increase depending on whether selection is homogenous or variable among leaves (Christensen and Peet, 1984; Dini-Andreote *et al.*, 2015; Meiners *et al.*, 2015). **(d)** Ecosystem or host function is anticipated to be unimodal or saturating over a successional gradient (van Ruijven and Berendse, 2005; Cardinale *et al.*, 2007; Weis *et al.*, 2007; Armitage, 2016), — a pattern predicted to be influenced by **(e)** the positive effects of *α*-diversity and living biomass on ecosystem function (Hooper *et al.*, 2005; Bell *et al.*, 2005). Dashed lines denote alternative hypotheses.

To test these hypotheses, I followed cohorts of pitcher leaves over three years and quantified their associated digestive communities through time. In addition, I measured these communities’ rates of decomposition, respiration, and their host leaves’ nitrogen uptake efficiencies. These data were used to test whether host-associated digestive communities follow general, predictable successional patterns and whether their turnover can influence a host’s ability to digest prey and sequester nutrients.

## MATERIALS AND METHODS

Complete documentation of the study system, data collection, and statistical analyses are provided in the supplementary materials and methods.

### *In situ* isotopic labeling of pitcher leaves

A stable isotope pulse-chase experiment was used to measure rates of decomposition and nitrogen cycling by the pitchers’ aquatic food webs. In early June 2013, I identified and tagged 50 unopened *Darlingtonia* pitcher leaves of equivalent age on different plants growing in a large population in the Plumas National Forest (Plumas Co., CA). Pitcher leaves remain sterile until completing their development and commencing prey capture, and each leaf has a lifespan of approximately 1.5 years. On the day the pitcher leaves first opened in mid-June, I fed gel capsules containing 20 sterile, ^15^N-enriched fruit flies (*Drosophila melanogaster*) to five random leaves, which were then left undisturbed for 11 days. I returned to the site to remove these ^15^N-labeled pitcher leaves and to feed isotope-labeled flies to 5 additional leaves belonging to the same cohort. This process was repeated every 11 days up to day 88 (mid-September), and again on day 365 (June 2014) with 10 leaves. Because the weight of enriched flies (4.25 mg) was much smaller than the average (177 mg) and standard deviations (176 mg) of natural prey masses within a leaf age class, this prey addition was unlikely to significantly overwhelm the natural variation in nutrient levels experienced by pitcher food webs. The 11-day timeframe was chosen based on preliminary data demonstrating peak N incorporation rates by lab-reared plants between 4 and 11 days after prey capture. I repeated this experiment in 2014-2015 in a nearby population of *D. californica* and included an additional 166-day sample. The sampled leaves were placed on ice and quickly returned to the lab.

### Quantification of pitcher leaf communities through time

From each freshly-collected leaf I removed 700 microlitres (μL) of fluid for DNA extraction using the PowerSoil microbial DNA isolation kit (MoBio Laboratories, Inc.) and stored the extractions at -80° C. Next, I dissected the pitcher leaves and categorized the state of fruit fly decomposition on an ordinal scale from 0 (no decomposition; flies undamaged) to 5 (completely decomposed; head capsules and wings only). I identified and enumerated all protists and living arthropods (primarily *Sarraceniopus* mites and *Metriocnemus* midge larvae) in each leaf’s fluid and interior surface under a light microscope and used epifluorescence microscopy to enumerate SYBR-Gold (Thermo Fisher Scientific, Inc.) stained bacterial cells and virus-like-particles bound to 0.02 micrometer (μm) filters. All prey detritus in a leaf was oven-dried at 60° C and weighed.

#### Bacterial community sequencing

Extracted DNA was sent for PCR amplification of the 16S SSU-rRNA genes (primer set 515f/806r) and multiplexed 2×150 bp paired-end sequencing on the Illumina MiSeq at the Argonne National Lab Core Sequencing Facility (Lemont, IL). Sequences were deposited on the MG-RAST public server (http://metagenomics.anl.gov/) server under project ID mgp14344. The QIIME bioinformatics pipeline was used to assemble and cluster reads into 97% operational taxonomic units (OTUs) (Caporaso *et al.*, 2010). I calculated each community’s alpha diversity (Shannon’s H, richness, phylogenetic) and beta diversity (Jensen-Shannon distance and weighted/unweighted UniFrac — a measure of community phylogenetic dissimilarity) using the *vegan* and *PhyloSeq* R packages, and used library size factor (LSF) normalization for all beta diversity metrics (McMurdie and Holmes, 2013; R Development Core Team, 2015; Oksanen *et al.*, 2015; Love *et al.*, 2014). Beta diversities for each sampling period were estimated using average inter-sample distances, and the results were unchanged when distances-to-centroid were used. I tested whether community composition changed with pitcher age using permutational analysis of variance (Anderson, 2001) on samples’ Jensen-Shannon distances (JSD) and UniFrac distances and visualized these results using PCoA plots. Results were unchanged when rarefaction was used for normalization.

To assess the generality of successional turnover in pitcher communities, I modeled OTU counts using a negative binomial generalized linear model (GLM) (Love *et al.*, 2014). Models were fit using empirical Bayes and OTUs experiencing significant log_2_-fold change among time points were identified using Wald *p-*values. I defined the ‘successional microbiome’ as the subset of OTUs experiencing a statistically significant (*α* = 0.01) ≥ 8-fold change in abundance between any two pitcher age classes and used these OTUs to construct an abundance-weighted heat map. The predictive accuracy of this subset of OTUs was assessed by training a random forest machine learning algorithm on OTU counts from the 2013 study population and using it to predict the age of samples from the independent 2014 study population. Model accuracy was evaluated using the coefficient of determination (*R*^2^) for predicted vs. observed ages along a 1:1 line. The entire bioinformatic/analytical pipeline is illustrated in figure S1.

#### Estimating microbial community traits

The Biolog GN2 microplate assay (Biolog Inc., Hayward, CA) was used to measure the carbon substrate use patterns of the microbial communities from an independent collection of 11, 55, and 365 day-old pitchers (10 from each age in 2014). These time-points were chosen to represent early, middle, and late-stage communities. Plates were inoculated in triplicate using the same dilute, filtered, starved communities described above, and incubated for 3 days at 25° C. I regressed substrate counts against leaf age using a negative binomial GLM to determine whether the number of metabolized substrates differed among leaf community age. To visualize differences in substrate profiles between age classes, I plotted samples onto principal coordinate (PCoA) axes based on their Jaccard distances.

I used ancestral genome reconstruction implemented by the PICRUSt software (Langille *et al.*, 2013) to predict the rRNA copy number and functional gene contents for the subset of OTUs in my samples present in the greengenes database (nearest sequenced taxon index = 0.071 ± 0.01 SEM). I estimated the mean weighted rRNA copy number of each pitcher sample (Nemergut *et al.*, 2015) and then evaluated their temporal turnover using ANOVA. Pitcher samples were then ordinated based on their predicted level 3 KEGG pathway relative abundances (Kanehisa *et al.*, 2016) using principal components analysis (PCA) and then hierarchically clustered. I filtered KEGG pathways using ANOVA *p*-values (*p* ≤ 0.01) and effect sizes (η^2^ ≥ 0.26) in order to identify genes and pathways (focusing primarily on enzymes involved in protein degradation and nitrogen transformation) that were predicted to be differentially enriched across time points. The predictive nature of these data precluded statistical hypothesis testing, and are treated as speculative hypotheses.

### Quantification of pitcher ecosystem properties through time

Empty pitcher leaves were thoroughly rinsed, dried at 60° C, homogenized in a bead-beater, weighed, and analyzed for ^15^N using an isotope ratio mass spectrometer at the UC Davis Stable Isotope Facility (Davis, CA). I used the fly and leaf ^15^N measurements to estimate the total amount of fly-derived ^15^N found in a leaf’s tissue after 11 days, which is interpreted to be the host leaf’s nitrogen uptake efficiency.

To estimate each pitcher microbial community’s potential C-respiration rate, I inoculated starved, washed pellets of pitcher bacteria into deep-well plates containing 800 μL sterile medium comprised of M9 salt solution and ground cricket powder. I used the MicroResp^TM^ respirometry system to measure the rates of CO_2_-C respired from cultures over three days at 25° C. These rates of CO_2_ respiration reflect the potential respiration rates of each pitcher’s bacterial community in a common environment.

I assessed temporal variation in pitcher ecosystem properties using ANOVA for N uptake efficiency/carbon respiration and a multinomial logit model for the fly decomposition category (Agresti, 2013). Covariates in these models included bacterial biomass and diversity, midge larvae abundance, leaf dry weight, and leaf age. Best-fit models were identified pluralistically using a combination of R^2^ and small-sample adjusted Akaike Information Criterion (AIC_c_) statistics (Burnham and Anderson, 2003). To investigate whether bacterial community composition influenced host functioning, I ran a Mantel test to assess whether pairwise Euclidean distances among samples’ N uptake efficiencies covaried with their pairwise JSD or UniFrac dissimilarity metrics.

### Verifying the effects of community structure on host function

Pitcher leaves of differing ages might physiologically regulate nitrogen uptake independent of their associated food webs, which can obscure food web effects. To account for this, I ran a field experiment to separate the effects of the food web and host leaf age on rates of N uptake. During late July 2014 I identified 15 pitcher leaves aged 11 days, 55 days, and > 365 days (5 leaves of each age), intended to represent young, middle-aged, and senescing pitchers based on developmental trends observed the previous year. The fluid from these leaves was removed and mixed in equal parts to form a homogenate. 5 mL aliquots of these homogenized communities were then returned to the host plants. Additionally, 20 ^15^N-enriched fruit flies were delivered into each leaf. I returned after 11 days to harvest and process these pitchers for N-uptake efficiency as previously described. I used ANOVA to test whether the N-uptake efficiencies of these pitchers with homogenized food webs recapitulated the N-uptake patterns from natural pitcher food webs of equivalent age from the same population.

## RESULTS

### Temporal changes in the *Darlingtonia* food web

The dynamics of dead and living biomass were qualitatively similar to the predictions in figure 1a. Pitcher leaves’ prey biomass varied widely among leaves of the same age, and mean prey masses quickly increased after opening and remained relatively stable throughout the plant’s lifespan (Figure 2a). Bacterial biomass also rapidly accumulated in young pitcher leaves and increased over time during the first growing season to a maximum of 1×10^11^ cells mL^-1^ before declining during the second growing season (Figure 2a). Virus-like particles, *Sarraceniopus darlingtonae* mites, and *Polytomella agilis* flagellates also increased in abundance during the first growing season (Figs. 2a, S1). In addition to *P. agilis,* I detected numerous other eukaryotes, including *Bodo*, *Monas*, *Petalomonas*, *Rhynchobodo*, *Chilomonas*, *Colpoda*, *Philodina*, and *Chlamydomonas*, but these taxa were observed in 10 or fewer pitcher leaves with no apparent temporal trends in occupancy or richness (Figure S2). Likewise, I did not detect a temporal trend in bacterivore beta diversity among time points until they diverged in year 2 (Figure S2).

**Figure 2.**
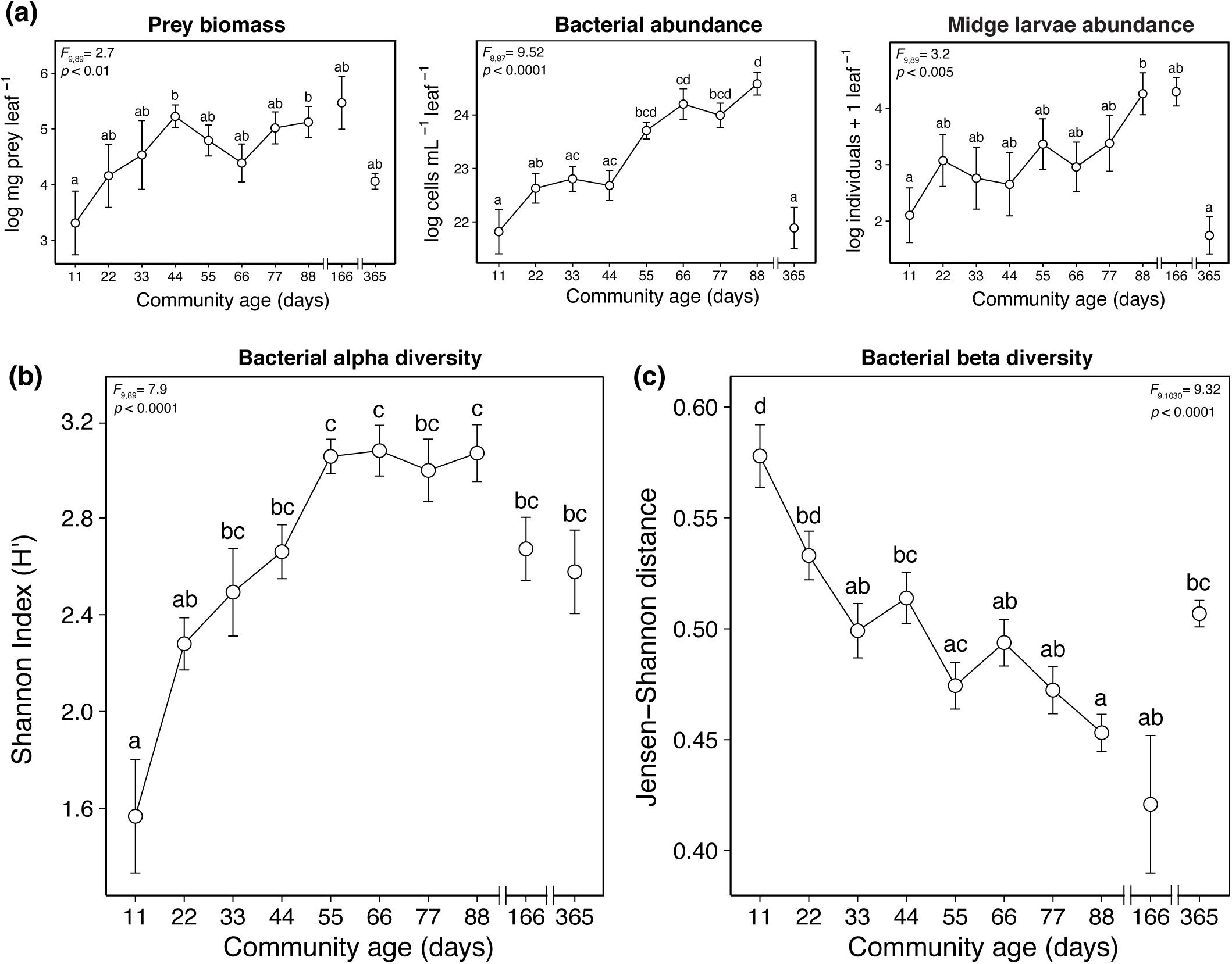
Trends in community composition during succession. **(a)** Insect prey biomass rapidly increased in leaves after opening and remained relatively steady throughout the remainder of the leaf’s lifespan, while bacterial and midge larval abundances steadily increased throughout leaves’ first growing season, and then sharply declined after the first year. **(b)** Bacterial alpha diversities increased and then leveled off in middle-aged pitcher communities, dropping slightly during year 2. **(c)** Conversely, leaf bacterial beta diversities decreased during the first growing season and increased at the beginning of year 2. In each graph, shared letters above groups indicate no significant pairwise differences (*p* > 0.05). Points denote mean values ± SEM.

### Composition and convergence of pitcher bacterial communities

After quality filtering of 16S amplicon sequences, the final OTU table consisted of 3 642 446 total reads representing 762 97% OTUs. The minimum and maximum number of reads per sample (n = 99) were 21 983 and 83 157, respectively (mean = 36 972), and read counts did not differ among age classes (*F*_9,89_ = 1.3, *p =* 0.26). Of the top 50 most abundant OTUs detected across pitcher samples, the majority belonged to families Bacteroidetes (Figure S3), Firmicutes (Figure S4), and Proteobacteria (Figure S5). As hypothesized in figure 1a, bacterial alpha diversities (Shannon’s H’) peaked at the end of the first growing season and experienced a slight decrease after day 88 (Figure 2b), whereas phylogenetic diversity increased over the entire study period (Figure S2). Taxonomic richness was highly correlated with phylogenetic diversity (Pearson’s *r* = 0.96) — increasing over time with the greatest variation among the 365-day samples. In contrast with the prediction in figure 1c, however, community composition tended to converge (i.e., beta diversity decreased) during the course of the first growing season, and diverge again during the start of the second growing season, according to both taxonomic (Figure 2c) and phylogenetic (Figure S2) dissimilarity metrics. Furthermore, permutational ANOVA on Jensen-Shannon and UniFrac distances revealed a structuring of pitcher bacterial communities by age class (table S1) and parallel successional trajectories between years (Figs. 3a, S6).

**Figure 3.**
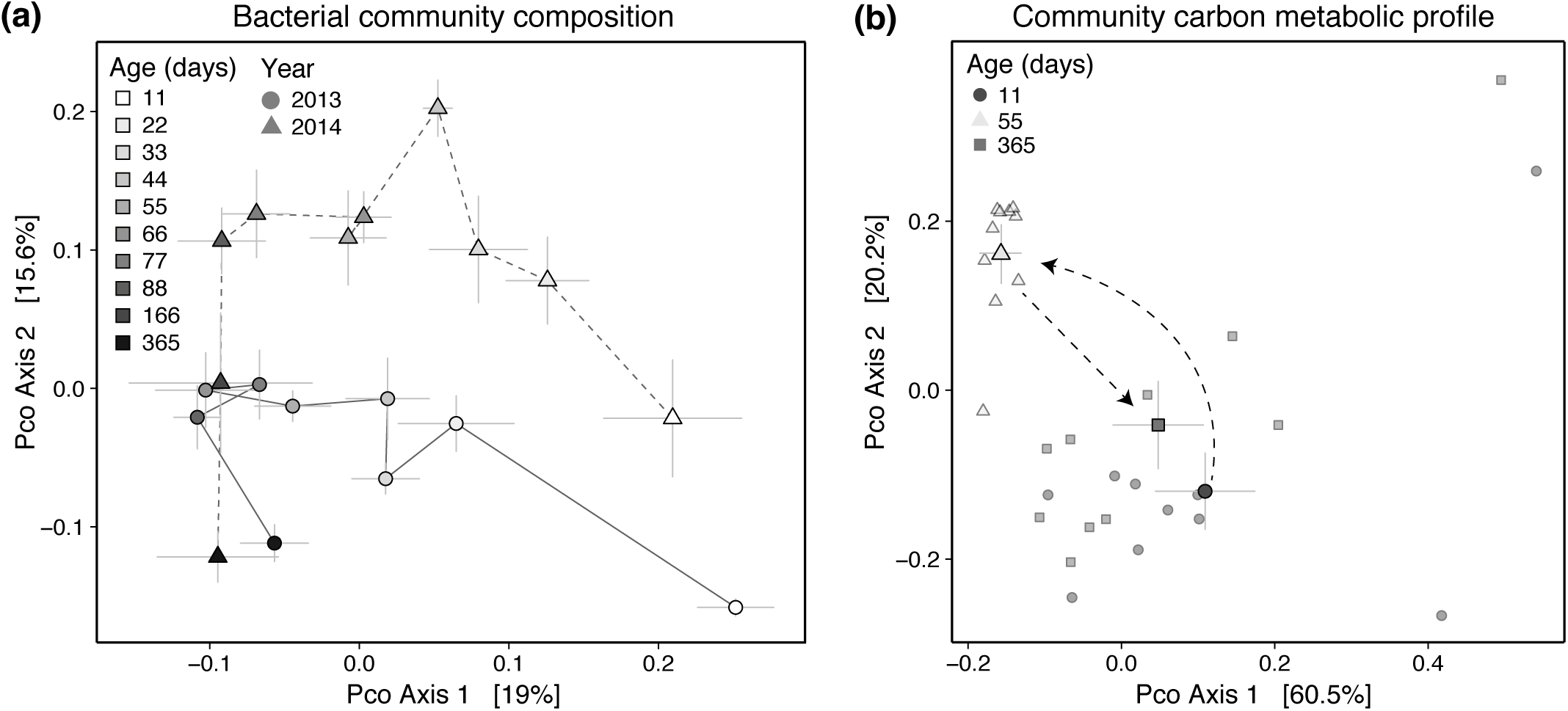
Principal coordinate (PCoA) plots for **(a)** Jensen-Shannon distances between samples, demonstrating convergence and approximately-parallel successional trajectories in between-population community structures over time, and **(b)** Jaccard distances between Biolog ^TM^ plates for communities of different ages, demonstrating the convergence of metabolic profiles in mid-successional pitcher leaves and overlapping metabolic profiles for young and senescing leaves. The percentages of variance explained by the principal coordinates are displayed on each axis. Points denote yearly centroid values ± SEM.

A subset of OTUs experienced particularly strong temporal turnover (Figure 4). These taxa fell primarily into the phyla Proteobacteria (37 OTUs), Bacteroidetes (16 OTUs) and Firmicutes (14 OTUs). Using these OTUs to train a random forest classifier to predict the pitcher community’s age resulted in a high classification accuracy for withheld data (observed vs. predicted *R*^2^ = 0.80). Likewise, a random forest trained on 2013 data was successful at predicting the ages of samples collected from the independent 2014 population (*R*^2^ = 0.75) (Figure S8), implying that observed community trajectories are parallel and generalizable between individuals and populations.

**Figure 4.**
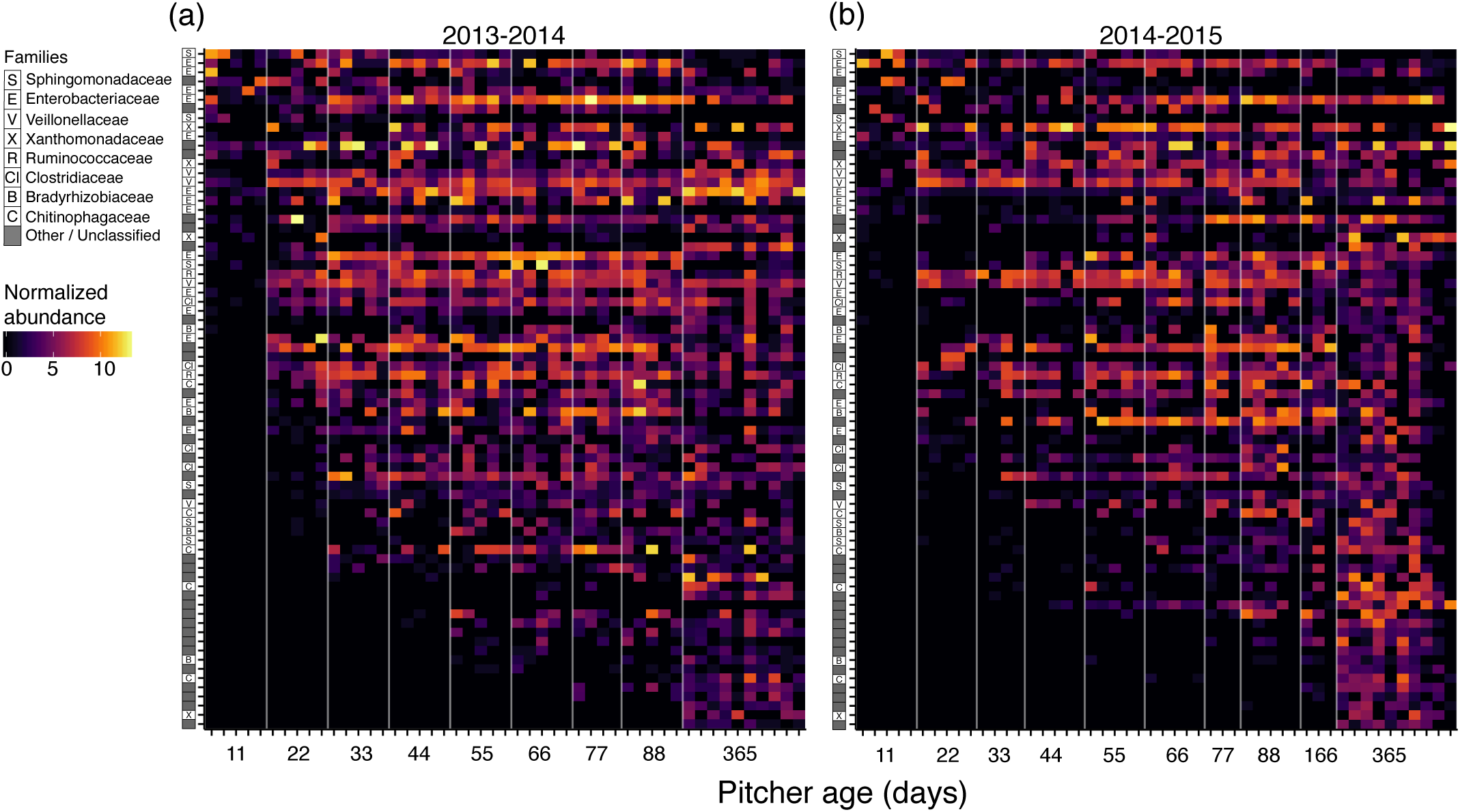
Abundance-weighted heat map of 97% OTUs that experienced significant (*p <* 0.01) 8-fold or greater turnover between time points for the **(a)** 2013 Blackhawk Creek and **(b)** 2014 Butterfly Valley study populations. Tick marks on X-axis denote individual pitcher samples. OTUs are labeled by family and ordered based on the community age in which they were first detected regardless of year. Random forest models trained on OTU abundances from 2013 were able to predict the ages of 2014 samples with 75% accuracy (Figure S7).

### Temporal trends in the functional attributes of pitcher microbiota

Assays of pitcher communities’ carbon substrate use patterns mirrored trends in taxonomic and phylogenetic alpha and beta-diversities — namely, early and late-stage pitcher communities both metabolized significantly fewer carbon substrates than did 55-day communities (Figure 5a). Furthermore, 11-day and 365-day pitchers’ substrate profiles were much more variable than and clustered apart from the 55-day samples. (Figs. 3b, 5b).

**Figure 5.**
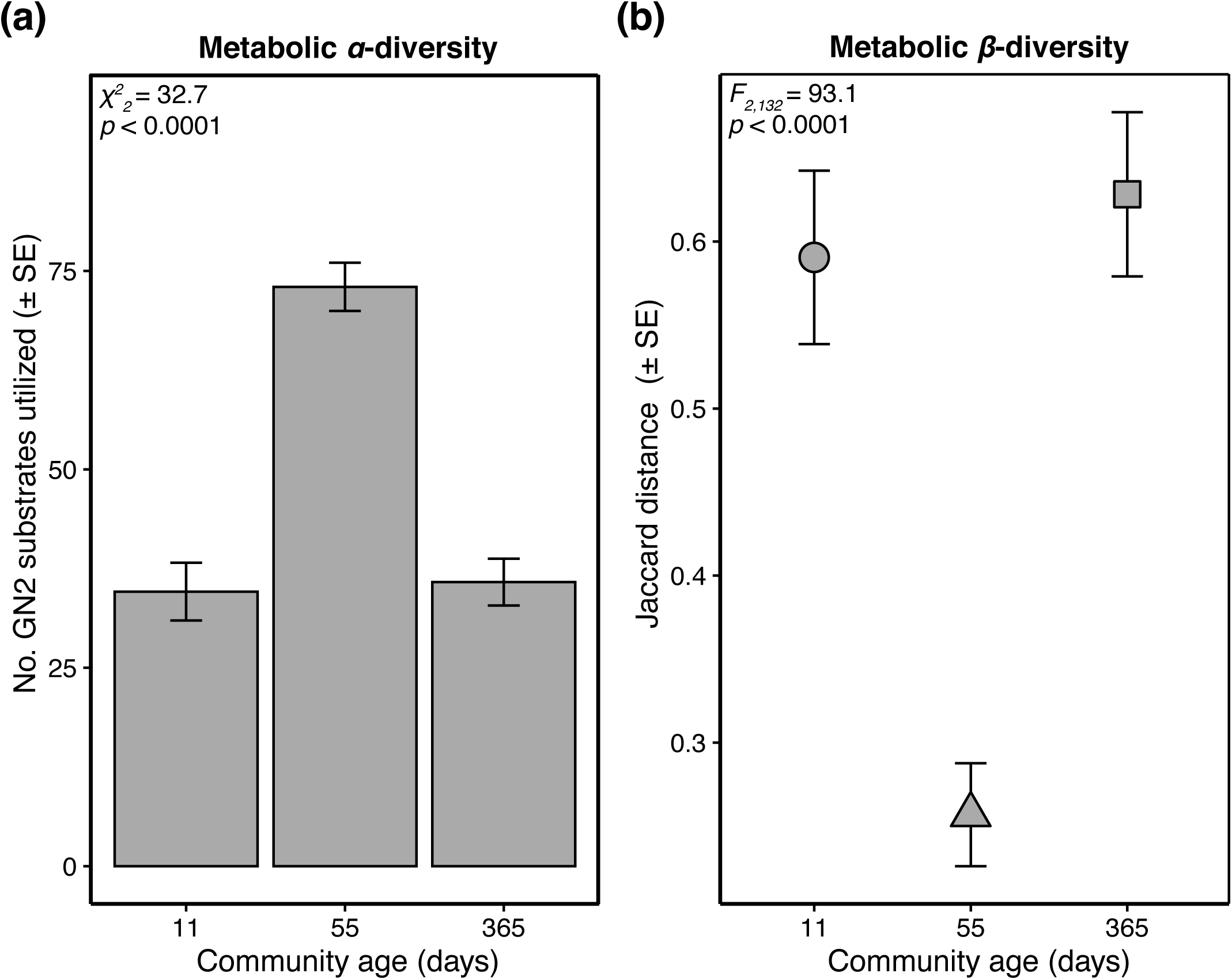
**(a)** Mid-successional pitcher communities were capable of metabolizing significantly more Biolog GN2 plate C-substrates than were early- and late-stage communities. **(b)** Mid-successional pitcher communities were much more similar to one another in terms of their carbon metabolic profiles than were early- and late-stage pitchers.

A PCA plot of samples’ reconstructed metagenomes predicted pitcher samples to separate by age, with the greatest distances between the 11-day and 365-day communities (Figure S8). The average number of rRNA gene copies per taxon was predicted to be greater in 11-day pitchers than in any other age class (Figure S9). This trend was also observed in the relative abundances of a number of other predicted KEGG pathways, such as flagellar assembly, motility, chemotaxis, and ABC transporters (Figure S10). Conversely, a variety of metabolic pathways were predicted to increase over time (Figure S11). Likewise, the abundances of genes involved in nitrogen cycling (deamination, nitrogen mineralization, denitrification, and nitrogen fixation) were also predicted to increase over a pitcher leaf’s lifespan (Figs. S12-S15).

### Linking community dynamics and ecosystem properties

Prey decomposition was unimodal over leaves’ lifespans, peaking at 44-88 days (Figure 6a). This increased decomposition, however, did not herald similar temporal differences in common-garden community respiration rates, although there was still a positive, non-significant unimodal trend in mean respiration rates over time (Figure S2). Multinomial logit models predicted bacterial diversity, bacterial abundance, and midge abundance to positively influence a pitcher’s probability of having a higher decomposition score (Figures 6b and 6c, Table 1). Leaf nitrogen uptake efficiency also increased during the first growing season and subsequently declined at the start of year 2 (Figure 6d), and was found to be positively associated with decomposition extent and leaf dry mass (Figure 6e, Table 1). Additionally, there was a weak but significant positive correlation between pitcher samples’ JSD/unweighted UniFrac distances and their Euclidean distances in nitrogen uptake efficiencies (JSD Mantel *r* = 0.08, *p* < 0.05; UniFrac Mantel *r =* 0.10, *p* < 0.05). Finally, in contrast to natural pitcher samples collected in 2014, the nitrogen uptake efficiencies of experimentally-homogenized pitcher food webs did not differ between leaf age classes (*F*_*2,12*_ = 0.98, *p* = 0.40) (Figure 7).

**Table 1.**
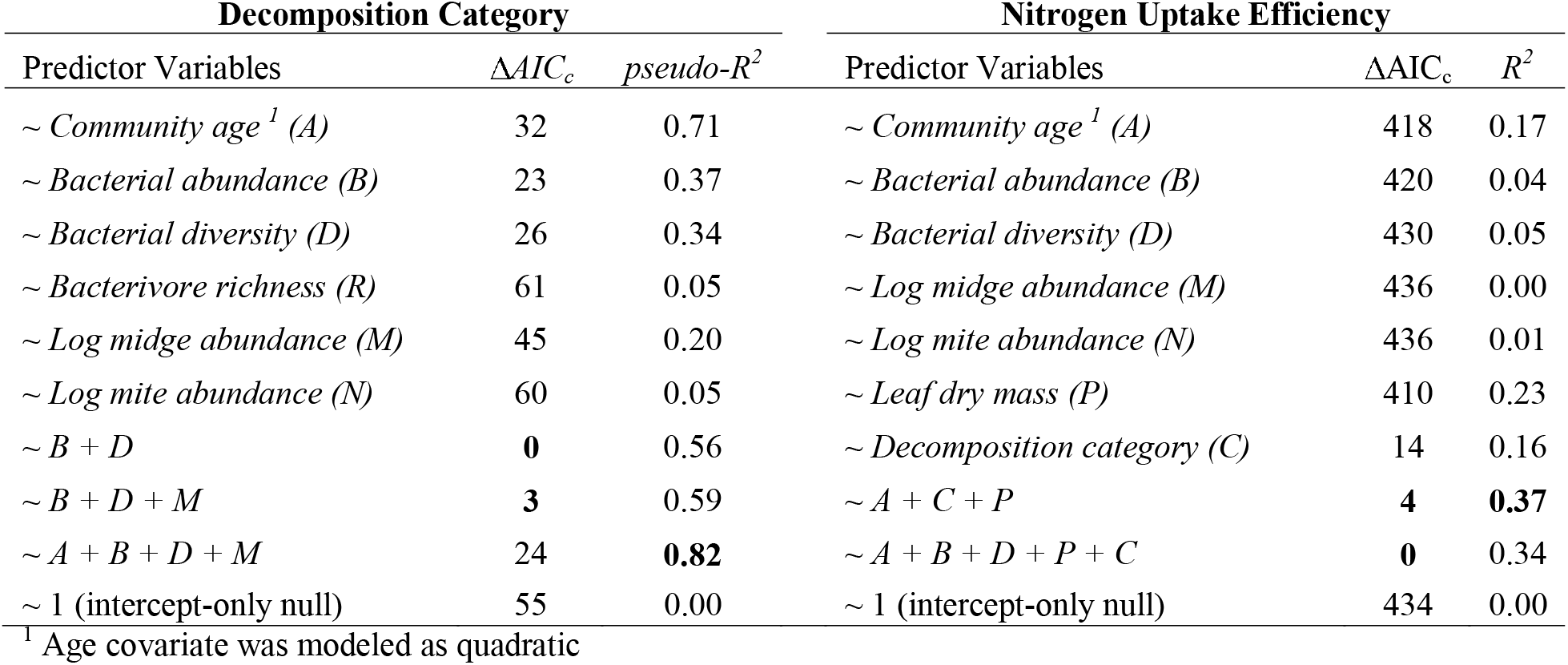
Model selection results of multinomial logit and linear regression models for decomposition category and nitrogen uptake efficiency, respectively. Bolded values indicate the best-performing models based on AIC_c_ and *R*^2^ statistics. AIC_c_ values falling within 9 units of the top model were considered equally parsimonious.

| Decomposition Category |  |  | Nitrogen Uptake Efficiency |  |  |
| --- | --- | --- | --- | --- | --- |
| Predictor Variables | $\Delta AIC_c$ | $pseudo-R^2$ | Predictor Variables | $\Delta AIC_c$ | $R^2$ |
| ~ Community age<sup>1</sup> (A) | 32 | 0.71 | ~ Community age<sup>1</sup> (A) | 418 | 0.17 |
| ~ Bacterial abundance (B) | 23 | 0.37 | ~ Bacterial abundance (B) | 420 | 0.04 |
| ~ Bacterial diversity (D) | 26 | 0.34 | ~ Bacterial diversity (D) | 430 | 0.05 |
| ~ Bacterivore richness (R) | 61 | 0.05 | ~ Log midge abundance (M) | 436 | 0.00 |
| ~ Log midge abundance (M) | 45 | 0.20 | ~ Log mite abundance (N) | 436 | 0.01 |
| ~ Log mite abundance (N) | 60 | 0.05 | ~ Leaf dry mass (P) | 410 | 0.23 |
| ~ B + D | **0** | 0.56 | ~ Decomposition category (C) | 14 | 0.16 |
| ~ B + D + M | **3** | 0.59 | ~ A + C + P | **4** | **0.37** |
| ~ A + B + D + M | 24 | **0.82** | ~ A + B + D + P + C | **0** | 0.34 |
| ~ 1 (intercept-only null) | 55 | 0.00 | ~ 1 (intercept-only null) | 434 | 0.00 |

<sup>1</sup> Age covariate was modeled as quadratic

**Figure 6.**
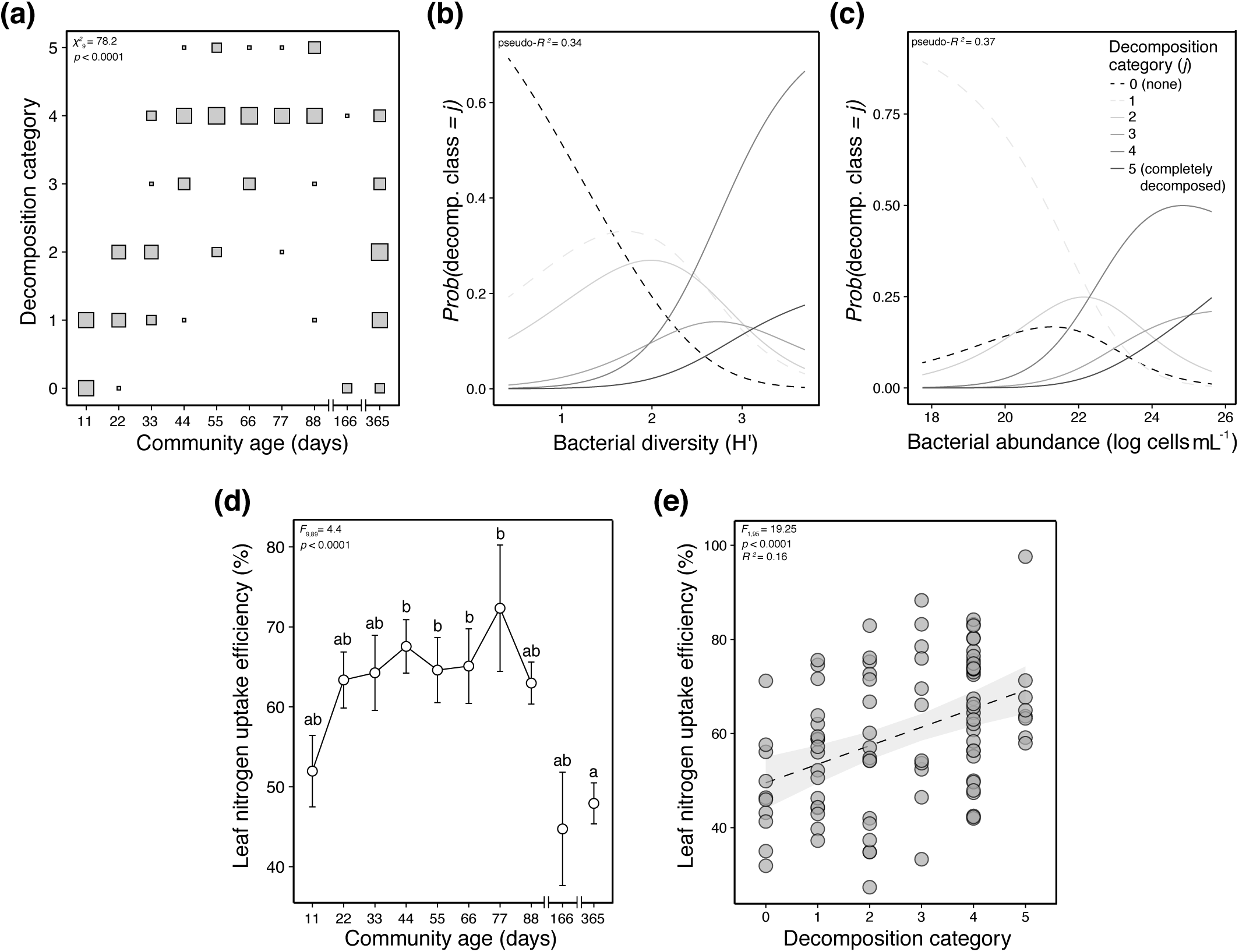
Trends in ecosystem properties during succession. **(a)** The frequencies of decomposition classes for pitchers of different ages. Square size is proportional to relative frequency of a particular decomposition category for that age class. χ^2^ is the likelihood ratio test statistic for the effect of pitcher age on the fit of a multinomial logit distribution to predict decomposition categories. **(b&c)** The probabilities of observing high decomposition rates increases with both bacterial diversity and bacterial biomass. Curves represent fitted proabilities of multinomial logit models, and individual curves can be interpreted as logistic regression fits for each decomposition category. **(d)** Pitcher leaves’ nitrogen uptake efficiencies change over time, and are significantly lower in late-stage pitcher leaves. Points denote mean values ± SEM. **(e)** The extent of prey decomposition is positively associated with the percentage of prey-derived nitrogen found in the host leaf’s foliar tissue. The dashed line denotes the best-fit linear model ± 95% CI.

**Figure 7.**
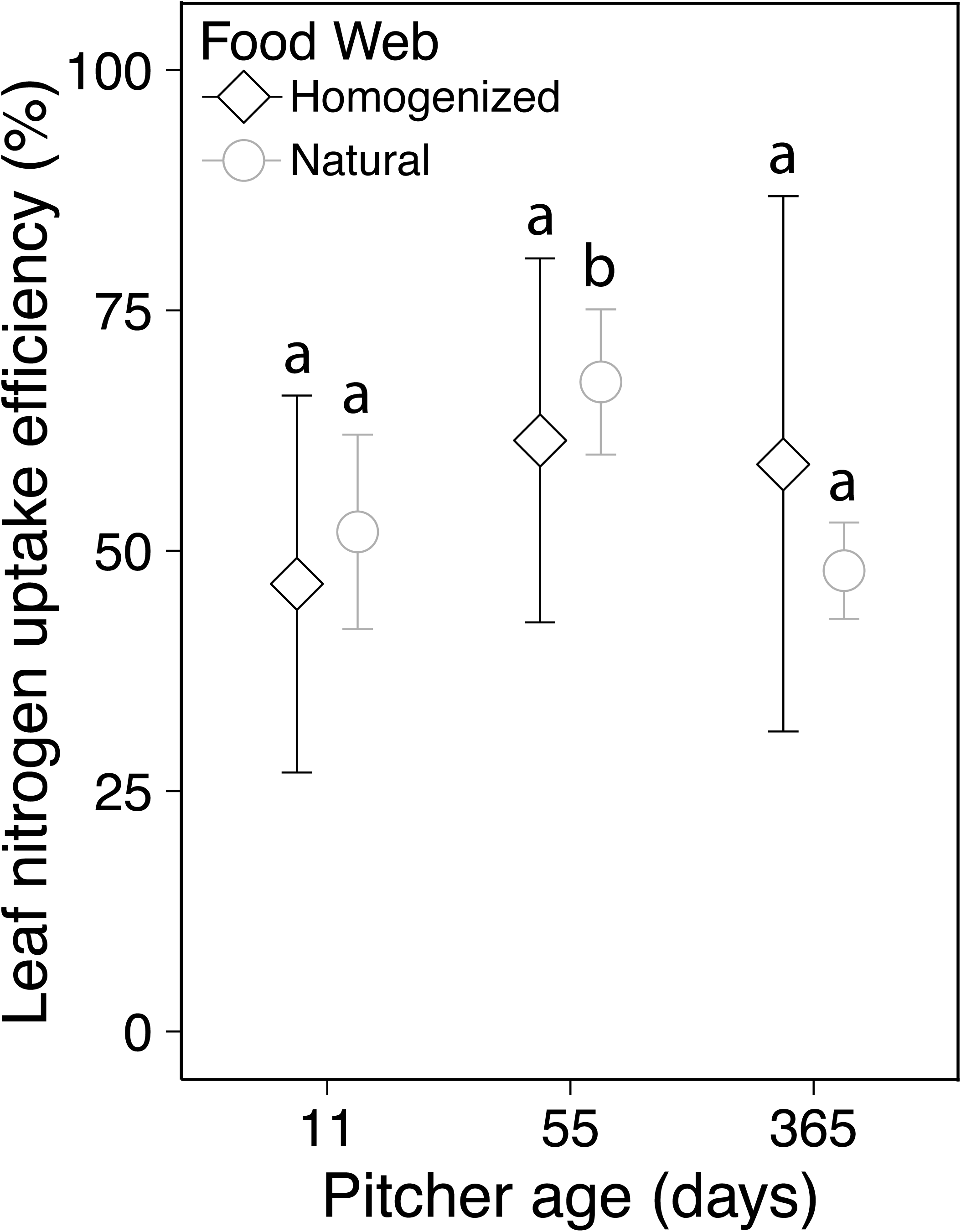
Homogenizing the food webs of 11, 55, and 365 day pitchers and placing them back into the plants removes the significant differences observed in natural pitcher communities of the same ages. Letters above the groups represent the within-treatment contrasts. Points denote mean values ± SEM (n = 5).

## DISCUSSION

As predicted, community diversity and biomass were positively associated with rates of prey decomposition, and the extent of decomposition was positively associated with the fraction of prey-derived nitrogen removed from the food web by the host leaf. In concert, these results imply that the services these digestive communities provide their hosts are time-dependent — highlighting important, general linkages between the temporal dynamics of communities and rates of ecosystem or host function.

### Temporal patterns in community composition

The logistic-like accumulation of both bacterial diversity and biomass in developing pitcher leaves aligns with both predictions from succession models (Figure 1b) and time series of animal gut communities (e.g., Koenig *et al.*, 2011; Jemielita *et al.*, 2014). However, it is important to note that both bacterial and midge abundances decreased over the winter — likely in response to the cessation of prey capture. A unimodal or monotonic increase in diversity over time is anticipated for open systems experiencing high rates of immigration and low rates of extinction, which is the likely state of pitcher leaves during their first growing season. Once leaves cease to produce prey attractants, prey capture becomes more stochastic (Wolfe, 1981). Because of this, bacterial communities may experience extinctions under diminishing resource levels or continue to accumulate diversity if prey capture continues to occur. This may explain the increased variation in diversity among year-old leaves.

Because pitcher plant leaves are similar in habitat structure and resource composition at a particular point in time, it is not surprising that bacterial communities converged in composition over the first growing season. This convergence can be attributed to common selection pressures acting on a shared pool of immigrants, which would serve to homogenize communities (see Vellend, 2016). This convergence is supported by the converging carbon substrate and OTU profiles of pitcher communities from two different populations. Successional convergence has also been documented in non-bacterial communities from the pitcher plant *Sarracenia purpurea* (Miller and terHorst, 2012), other phyllosphere bacterial communities (Copeland *et al.*, 2015), the human gut (Palmer *et al.*, 2007), and more generally, across a variety of terrestrial (e.g., Christensen and Peet, 1984) and aquatic ecosystems (e.g., Moorhead *et al.*, 1998).

Contrasting with this pattern, year-old leaves contained higher microbial beta diversities than those observed in preceding time points, implying communities diverged over the winter. This is likely the consequence of stochastic prey capture amplifying differences in leaves’ ratios of labile to recalcitrant metabolic substrates. If this ratio constitutes a reasonably strong selection gradient, then this heterogeneity should drive divergence among communities (Eisenhauer *et al.*, 2013; Dini-Andreote *et al.*, 2015). Alternatively, stochastic drift can drive community divergence when the number of individuals is small (Orrock and Watling, 2010; Vellend, 2016). In *Darlingtonia* leaves, however, drift is likely minimal, since bacterial population sizes are probably too large to be influenced by demographic stochasticity.

Temporal variation in propagule supply can also lead to community divergence (Evans *et al.*, 2017). This might occur when a fraction of ageing leaves, whose communities had previously been homogenized by a sustained input from a common microbial pool, suddenly experience a more stochastic supply of immigrants. If the species comprising the common immigrant pool also vary over time, then discontinuous, stochastic prey input could drive divergence in communities in the absence of drift and selection effects. A 55-year study of old-field communities observed similar patterns of convergence giving way to divergence driven by dispersal limitation (Meiners *et al.*, 2015). Nonlinear temporal trends in beta diversity have also been identified in host-associated and groundwater microbial communities (e.g., Marino *et al.*, 2014; Zhou *et al.*, 2014), though the processes governing these patterns remain vague. Pitcher microbial communities offer a tractable system in which to experimentally assess the relative influences of deterministic vs. stochastic dispersal on beta diversity.

### Temporal trends in communities’ functional attributes

Leaf communities’ carbon metabolic profiles had temporal patterns similar to OTU beta diversity, implicating a link between community composition and metabolic functioning. However, microbial community sequences were not generated from the leaves used for Biolog assays, prohibiting a direct test of this hypothesis. Many of the genes predicted to be enriched in young pitchers (ribosomal RNA copy number, chemotaxis/motility genes) have been linked to a taxon’s responsiveness to unpredictable nutrient conditions (Klappenbach *et al.*, 2000; Livermore *et al.*, 2014; Nemergut *et al.*, 2015). These predictions are in accordance with successional tolerance and inhibition models, wherein ruderal, fast-responders are eventually joined or outcompeted by more growth-efficient forms (Connell and Slatyer, 1977; Huston and Smith, 1987; Tilman, 1990).

Metabolic pathways contributing to amino acid demamination and N mineralization were predicted to be enriched during mid-succession — a pattern also detected during microbial succession on decomposing corpses (Metcalf *et al.*, 2016). Similar successional increases in metabolic genes have been documented in host-associated (Koenig *et al.*, 2011) and aquatic (Teeling *et al.*, 2012) bacterial communities. In concert with the community metabolic assays, these findings demonstrate, in principle, how bacterial communities’ taxonomic and functional profiles can undergo predictable changes over a host’s lifespan in accordance with predictions derived from succession models. The next step is to relate these community changes to the services they provide the host organism.

### Linking community properties to host functioning

In agreement with succession hypotheses (Figure 1d), detrital processing rates by the pitcher leaf communities varied over time, and were positively associated with detritivore abundances (bacteria, midge larvae) and bacterial diversity. Loreau (2001) reasoned that microbial diversity would enhance decomposition only if the number of organic compounds able to be metabolized by the community increased with alpha diversity. This prediction is supported by observations of peak bacterial diversity coinciding with peak carbon metabolic diversity during mid-succession (ca. 55 days). To date, the few studies to investigate microbial diversity and decomposition rates *in situ* have arrived at conflicting results (Hättenschwiler *et al.*, 2011) but a positive relationship is common in the few experimental tests using bacteria (Nielsen *et al.*, 2011), including in a lab experiment using bacterial isolates from the same *Darlingtonia* population studied here (Armitage, 2016). More generally, microbial community composition is anticipated to set ecosystem process rates (Figure 1e). — especially when the effects of environmental variation are minimal (Graham *et al.*, 2016).

From a host plant’s perspective, decomposition by its commensal biota should set limits on its rate of N sequestration. In *Darlingtonia*, the state of fly digestion explained a some of the variance in N uptake efficiency, though there was still a large amount of unexplained variance to account for. More convincingly, a follow-up experiment failed to detect the same mid-succession peak in N uptake efficiencies among pitcher leaves containing experimentally homogenized bacterial communities. The related pitcher plant *Sarracenia purpurea* also relies heavily on its bacterial community for nitrogen processing (Butler *et al.*, 2008). Furthermore, microbial community composition is an important determinant of nitrogen mineralization rates in soil (Balser and Firestone, 2005; Strickland *et al.*, 2009), and changes in N mineralization can track microbial community change over time, independent of environmental variation (Balser and Firestone, 2005). In concert, these results highlight the potential for the pitcher microbial communities to mediate N transfer from prey to host — a function critical to the fitness of a host plant adapted to life in nitrogen-poor soils.

Contrary to predictions from succession models (Vitousek and Reiners, 1975; Finn, 1982; Loreau, 1998), maximal rates of N loss from the *Darlingtonia* food web occurred during periods of high (rather than low) standing biomass. This mismatch may be explained by differences between donor-controlled food webs, which receive pulses of bioavailable N at regular intervals, and primary producer-controlled food webs, in which the N pool is slowly renewed *in situ* and quickly immobilized (Fierer *et al.*, 2010). As a consequence, donor-controlled food webs may not experience strong competitive pressure to sequester growth-limiting nutrients. This may be particularly true in *Darlingtonia* and other digestive communities for two reasons. First, rapid bacterial turnover (e.g., via viral lysis & protozoan grazing) serves to increase the concentration of bioavailable N. Second, pitcher leaves’ continuous accumulation of low C:N detritus (relative to plant-based food webs) may buffer the food web from a loss of nitrogen to the host plant.

### Succession or seasonality?

Because study leaves belonged to the same cohort, their temporal dynamics may reflect the effects of seasonal forcing rather than succession. Although winter temperatures drive the plants into a state of dormancy, their leaves persist, and I have observed active populations of mites, midges, and bacteria in pitcher leaves underneath snow cover, suggesting that the food web still functions during the winter months. Furthermore, these brief cold periods are unlikely to have caused strong population bottlenecks or extinctions, given the large bacterial biomasses observed across pitcher leaves. Seasonal forcing should cause community composition to be cyclical over an annual cycle, yet communities collected from 11 and 365-day samples on the same day were strongly dissimilar, implying that under nearly identical external environmental conditions, communities show measurable age-related differences — an observation in line with previous studies (Thompson *et al.*, 1993; Redford and Fierer, 2009; Williams *et al.*, 2013; Metcalf *et al.*, 2016).

### Cross-system considerations

It is now recognized that community and ecosystem dynamics are shaped by unique combinations of disturbances, competition, and dispersal (Meiners *et al.*, 2015). And although succession is most frequently defined in terms of species turnover, it is reasonable to redefine it as the change in average trait values or gene frequencies within a community. Such change could influence the functioning of the host plant if, for instance, selection favored a more efficient processing or storage of nitrogen by commensal organisms. The potential for rapid evolutionary change to influence ecosystem properties has been documented (Harmon *et al.*, 2009), yet theory integrating ecosystem development and evolution is scarce (Loreau, 1998). In doing so, care must be taken to avoid ascribing adaptive properties to ecosystems (i.e., treating ecosystems as ‘super-organisms’) (Odum, 1969). However, because many host-associated systems serve functions critical to their hosts’ fitnesses, they may be expected to more closely align with Odum’s controversial predictions for increasing stability and productivity. Tests of these predictions (e.g., Beaver, 1985; Neutel *et al.*, 2007) using existing quantitative frameworks (Finn, 1982; DeAngelis, 1992; Loreau, 1998) would be difficult but valuable contributions toward a unified theory of communities and ecosystems.

### Conclusions

By combining a ^15^N stable isotope pulse-chase experiment with observations of community dynamics, I have confirmed a number of successional hypotheses in natural, host-associated microbial digestive communities. In particular, my data support and extend the hypotheses of parallel community trajectories and mid-successional peaks in functional and taxonomic diversity to host-associated bacterial communities. In concert, these results represent a step towards integrating host-associated microbial communities into classical conceptual models of ecosystem development and demonstrate a coupling of community dynamics and host functioning. Looking ahead, more theoretical and experimental work is needed before we can identify definitive links between community dynamics and host functioning, and I believe that the continued experimental use of replicated, natural host-associated communities offers a productive path forward.

## ACKNOWLEDGEMENTS

I thank Anna Petrosky, Ramon Leon, & Stefani Brandt for assistance with data collection. Ellen Simms, Todd Dawson, and the UC Berkeley Forestry Camp provided facilities and equipment. I thank Stuart Jones, Mary Firestone, Mary Power & Wayne Sousa for critical feedback. Field collection permits were provided by Jim Belsher-Howe (USFS).

